# Spatial/molecular heterogeneity and treatment response in HER2+ early breast cancer: Translational analyses from the DAPHNe trial

**DOI:** 10.64898/2026.09.17.752407

**Authors:** Emily Chen, Ilana Schlam, Kenichi Shimada, Tianyu Li, Esther Ritah Ogayo, Ashka Patel, Janae Davis, Jia-Ren Lin, Austin Schultz, Madeline G. Townsend, Kelly Zheng, Carlos W. Wanderley, Ricardo Pastorello, Felicia N. New, Evelyn R. Metzger, Natalie Sinclair, Laura M. Spring, Meredith Faggen, Michael Constantine, Neelam Desai, Nadine Tung, Nabihah Tayob, Ian E. Krop, Sara M. Tolaney, Eric P. Winer, Jennifer Guerriero, Elizabeth A. Mittendorf, Adrienne G. Waks

## Abstract

Given the heterogeneity of HER2-positive breast cancer, reliable biomarkers to guide treatment are needed. We conducted multi-platform biomarker analyses of baseline tumor tissue from the DAPHNe trial of neoadjuvant paclitaxel/trastuzumab/pertuzumab (THP) for HER2-positive early breast cancer to characterize inter- and intra-tumor heterogeneity and to identify molecular predictors of response. A total of 98 patients with stage II-III HER2-positive breast cancer received neoadjuvant THP followed by surgery. Gene expression profiling, spatial protein profiling, and single-cell imaging (cyclic immunofluorescence) were performed on pre-treatment biopsies and a subset of residual disease specimens. Pathologic response was assessed using the residual cancer burden (RCB) score. Among HER2-positive patients included in the biomarker analysis, 34% of patients had node-positive breast cancer, and 66% had hormone receptor (HR)- positive tumors. HR-positive and HR-negative tumors differed significantly across gene expression, protein expression, and single-cell profiling. High ERBB2 gene signature, p53 gene signature, and HER2 protein expression predicted favorable response (RCB 0/1), while ESR1/PGR gene, estrogen receptor (ER) signaling gene signature, and ER alpha protein expression were associated with unfavorable response (RCB 2/3). Single-cell spatial analysis revealed that cancer cells clustered by shared HER2, ER, and PR expression, suggesting local expansion of phenotypically distinct subpopulations. ER-positive cancer cells were associated with lower HLA-A and PD-L1 expression, suggesting a less immunologically active tumor cell state. Baseline HER2 and ER expression are key predictors of response to neoadjuvant HER2-targeted therapy. Single-cell spatial profiling highlights intra-tumoral heterogeneity and suggests ER-driven immune exclusion as a potential mechanism of resistance.

## INTRODUCTION

Outcomes for patients with human epidermal growth factor receptor 2 (HER2)-positive breast cancer have greatly improved in the last few decades due to the development of HER2-directed therapies (*1-3*). Specifically, the standard of care for stage II or III HER2-positive breast cancer has evolved to entail neoadjuvant (pre-operative) chemotherapy with dual HER2-directed therapy (trastuzumab [H] and pertuzumab [P]) (*4*). However, it has become increasingly clear that HER2-positive breast cancer is a heterogeneous disease, with certain patients likely being overtreated by the current standard of care, and others recurring in spite of it (*5, 6*).

Thus, the focus of international research efforts in HER2-positive breast cancer has shifted toward more individualized or novel treatment strategies, including the abbreviation of neoadjuvant polychemotherapy to single-agent chemotherapy for some patients (e.g. in the CompassHER2-pCR, HELEN-006, and neoCARHP trials) (*7-9*), use of trastuzumab deruxtecan (T-DXd) in the neoadjuvant setting in the DESTINY-Breast11 trial [NCT05113251]) (*10*), and the use of neoadjuvant doublet HER2 blockade alone (without cytotoxic chemotherapy) in highly selected patients in the PHERGain trial (*11*). As results of ongoing clinical trials mature, the range of therapeutic options for patients with early-stage HER2-positive breast cancer is expected to expand. This underscores the growing importance of identifying reliable predictive biomarkers in the neoadjuvant setting, enabling personalized treatment strategies. Gene and protein expression profiling offer a powerful approach to elucidate tumor biology and uncover potential prognostic and predictive biomarkers of therapeutic response (*12-14*). Additionally, characterizing the tumor immune microenvironment (TME) can provide critical insights into prognosis, and may refine treatment decision-making (*13, 15*).

DAPHNe was a single-arm phase 2 trial that investigated the feasibility of abbreviating neoadjuvant treatment in patients with stage II-III HER2-positive breast cancer (*16, 17*). All patients received neoadjuvant paclitaxel-HP (THP), followed by adjuvant treatment determined by pathologic complete response (pCR) status. In this work, we leveraged a multi-platform profiling strategy to characterize both inter-tumoral heterogeneity (differences between tumors across patients) and intra-tumoral heterogeneity (cell-to-cell variability within individual tumors) on baseline and post-THP breast tumors. Bulk gene expression profiling and multiplex protein expression analyses were used to define inter-tumoral differences in intrinsic subtype, signaling programs, and immune context across the cohort, while single-cell spatial imaging enabled direct assessment of intra-tumoral heterogeneity in tumor cell states and their local microenvironment. Together, these complementary approaches offer a multi-dimensional view of HER2-positive breast cancer heterogeneity, with the potential to inform more individualized treatment strategies.

## RESULTS

### Patient and tumor characteristics

**Table 1** shows patient and tumor characteristics for the gene and protein expression biomarker populations, in comparison to the original trial population; all populations had similar characteristics in terms of patient demographics and clinicopathologic features of the tumors. Median patient age was 50 years, 34% of patients had node-positive breast cancer, 66% had hormone receptor (HR)-positive breast cancer, and 83% of tumors were HER2 immunohistochemistry (IHC) 3+ (with all patients being HER2-positive by national guidelines). Intrinsic subtyping of the cohort by PAM50 gene signature showed that 65% of the tumors were HER2-enriched, 16% were luminal A, 15% were luminal B, and 5% were basal-like. Overall, 56% of patients experienced pCR following neoadjuvant THP.

**Table 1:**
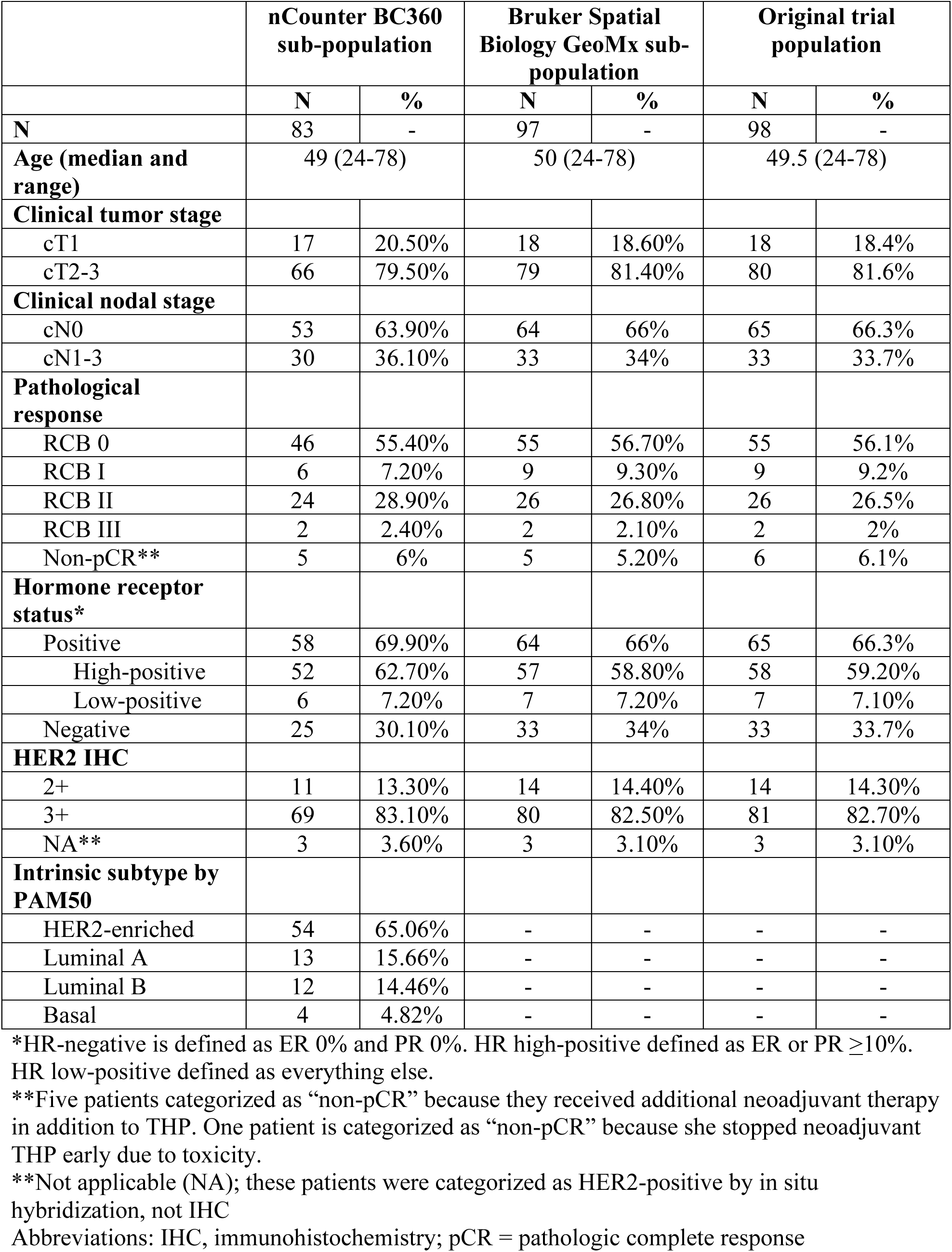
Patient and tumor characteristics.

### Gene and protein expression landscape of treatment-naïve HER2-positive breast cancer

To investigate molecular differences across clinical HER2 IHC score categories, we profiled 83 treatment-naïve breast tumors using the nCounter BC360 gene expression profiling panel (**Fig. S1A**). As expected, *ERBB2* expression was significantly higher in the HER2 IHC 3+ population compared to the IHC 2+ population (FDR p <0.05; **Fig. 1A**). No other significant gene expression differences were observed by HER2 IHC scores or nodal status (**Fig. S2**). There were multiple significant differences between HR-positive and HR-negative tumors. Compared with HR-negative tumors, HR-positive tumors exhibited significantly higher expression of *ESR1*, *PGR* and the ER signaling pathway, representing the three most differentially expressed genes/gene signatures (FDR-adjusted p <0.01). *PTEN* expression and the mast cells and apoptosis signatures were also significantly elevated in HR-positive tumors, albeit with relatively low fold-change. Among HR-negative tumors, two gene signatures were significantly overexpressed (FDR-adjusted p<0.05) relative to HR-positive tumors: BC p53 (a proxy for mutant-like versus wildtype-like p53 status in breast tumors, where a higher expression level corresponds to a more mutant-like phenotype) and homologous recombination deficiency (HRD) (**Fig. 1B**).

**Fig. 1:**
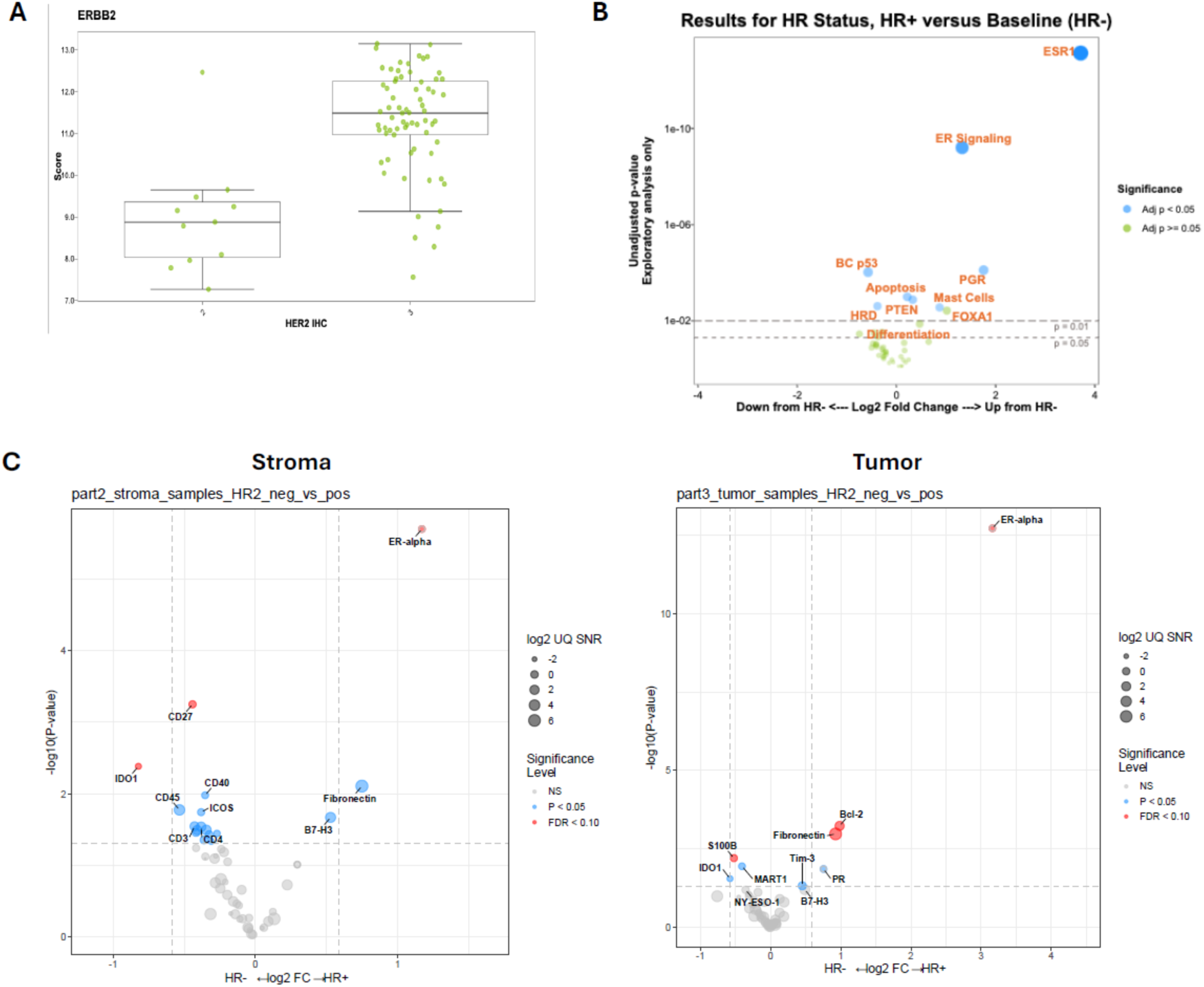
Gene and protein expression landscape of untreated HER2+ breast tumors. Bulk RNA was analyzed using the nCounter platform to test baseline single-gene and gene signature expression patterns across biological subgroups. **(A)** *ERBB2* single gene expression in tumors with clinically assessed HER2 IHC 3+ vs 2+ (FDR p-value <0.05 for difference). **(B)** Gene expression differences by HR+ vs HR-status. **(C)** Protein expression differences by HR+ vs HR-status. Abbreviations: ER, estrogen receptor; TIS, tumor inflammation signature.

At the protein level (**Fig. 1C; Table S1**), ER alpha was the most highly enriched protein in the tumor compartment of HR-positive compared with HR-negative tumors (fold-change 8.94, FDR adjusted p<0.05), consistent with both gene expression data and clinical definition of HR status. Expression of BCL-2, a regulator that promotes cellular survival by inhibiting pro-apoptotic proteins, was also higher in HR-positive tumors (fold-change 1.98, FDR adjusted p<0.05), as was expression of the extracellular matrix protein fibronectin (fold-change 1.90, FDR adjusted p<0.05). These features are consistent with a more hormonally driven, survival-associated tumor cell state in HR-positive disease. In contrast, the stromal compartment of HR-negative tumors exhibited higher expression of several immune-related proteins, including the immune checkpoint molecule PD-L1 (**Table S1)**, the co-stimulatory molecules CD27, CD40, ICOS, and 4-1BB, as well as the T cell markers CD3 and CD4 and CD45. Among these, only CD27 remained significant after multiple-testing correction (FDR adjusted p < 0.05), whereas others were significant only in unadjusted analyses (p < 0.05). Collectively, these findings suggest a more immune-enriched stromal microenvironment in HR-negative tumors.

### Gene and protein expression predictors of response to neoadjuvant THP

For all patients, response to neoadjuvant therapy was measured by RCB score at surgery, and responses were characterized as favorable (RCB 0/1) or unfavorable (RCB 2/3). We investigated baseline gene and protein expression features that correlated with response. In the gene expression data (**Fig. 2A, B**), *ERBB2* expression and the BC p53 signature were the significant predictors of favorable response, while *ESR1*, *PGR*, and the ER signaling signature were the strongest predictors of unfavorable response. These findings were mirrored by protein expression analyses (**Fig. 2C**; **Table S2**), which found that higher HER2 protein expression was the only significant predictor of favorable RCB response (fold-change 3.75, adjusted p<0.05) and ER alpha expression was the only significant predictor of unfavorable RCB response (fold-change 2.44, adjusted p<0.05). Among HR-positive/HER2-positive patients, higher HER2 protein expression in the tumor compartment remained the key predictor of favorable response to neoadjuvant THP among patients with HR-positive/HER2-positive tumors (fold-change 4.21, adjusted p<0.05; **Table S2**).

**Fig. 2:**
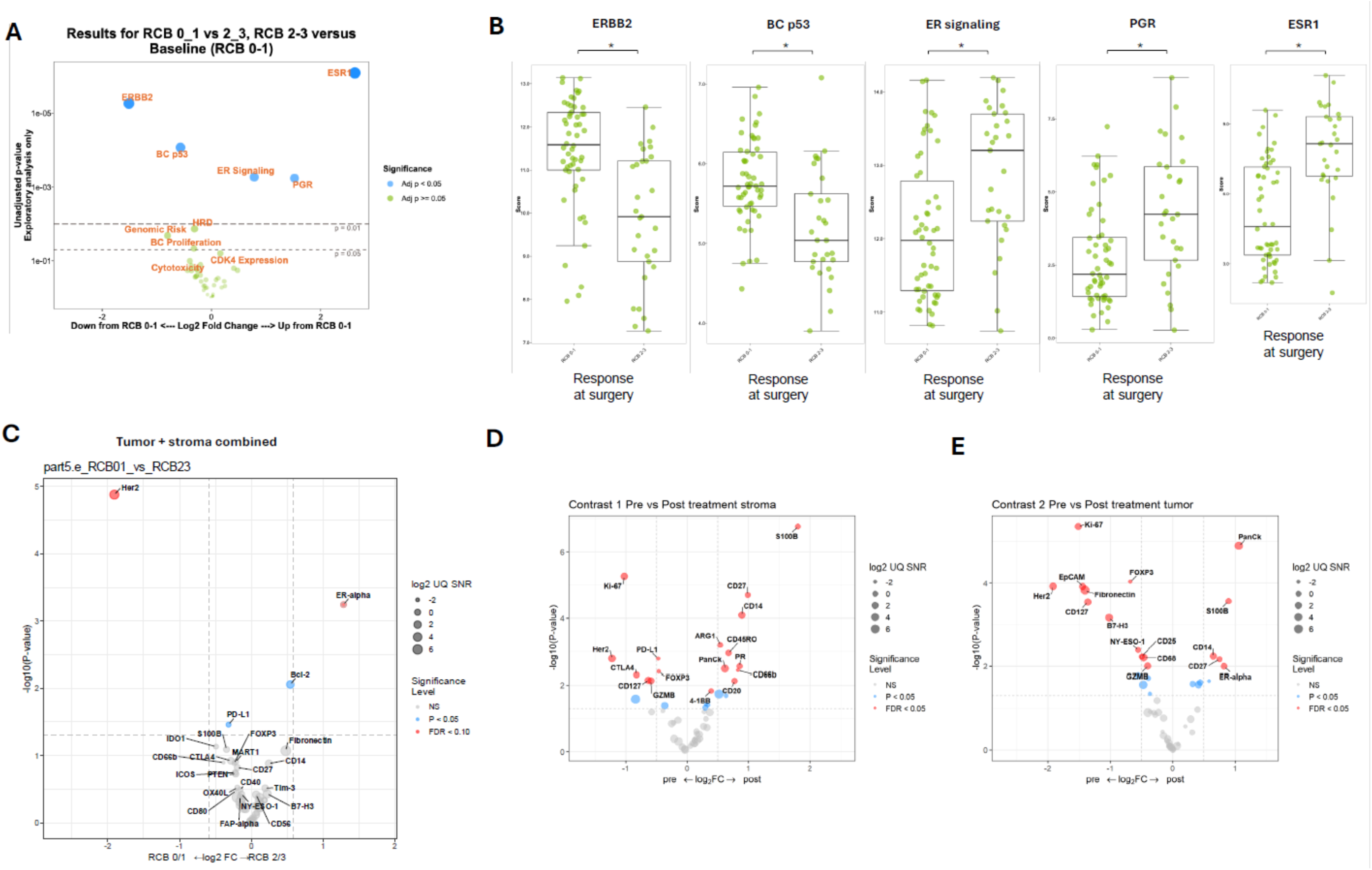
Gene and protein expression predictors of response, and evolution during neoadjuvant therapy. **(A)** Gene expression predictors of favorable (RCB 0/1) or unfavorable (RCB 2/3) response. X-axis indicates fold-change, with findings to the right of the panel being more likely in RCB 2/3 non-responders, and findings on the left of the figure being more likely in RCB0/1 responders. Y-axis indicates p-value. **(B)** For the five genes/gene signatures with significant differences between favorable and unfavorable responders in panel 2A, individual gene expression scores (y-axis) are shown. Asterisk indicate FDR-adjusted p-value for comparisons between subgroups. **(C)** Protein expression predictors of favorable (RCB 0/1) vs unfavorable (RCB 2/3) response. **(D)** Protein expression changes from pre- to post-treatment in the stromal compartment. X-axis indicates pre- vs post-treatment specimens. Y-axis indicates p-value. Presence of tumor cell-derived proteins eg Ki67, HER2, panCK, and PR likely represents contamination from the tumor cell compartment. **(E)** Protein expression changes from pre- to post-treatment in the tumor compartment. X-axis indicates pre- vs post-treatment specimens. Y-axis indicates p-value. Abbreviations: HR: hormone receptor, RCB: residual cancer burden.

### Evolution of features after neoadjuvant therapy

To assess the evolution of the tumor and microenvironment during neoadjuvant THP, we compared protein expression in pre-treatment breast core biopsies versus paired post-treatment breast surgical specimens for patients with significant residual disease at surgery (RCB 2/3). Of the 28 tumors included in this analysis (**Fig. S1B**), 26 were clinically defined as ER-positive, and 2 as ER-negative at baseline. In the stroma, expression of several proteins was higher at the pre-treatment timepoint (i.e., decreased during treatment), including immune checkpoint proteins PD-L1 and CTLA4, regulatory T cell marker FOXP3, T cell marker CD127, and cytotoxic effector molecule granzyme B (GZMB), whereas proteins with significantly higher expression in post-treatment samples included the pro-inflammatory calcium-binding protein S100B, T cell co-stimulatory proteins CD27 and 4-1BB, and granulocyte marker CD66b (**Fig. 2D**). These changes indicate complex remodeling of the stromal immune compartment during therapy. In the tumor cell compartment, HER2 and Ki-67 were significantly decreased following treatment, consistent with preferential elimination of highly HER2-driven and proliferative tumor cells. Expression of the epithelial marker EpCAM and the extracellular matrix protein fibronectin was also reduced, suggesting treatment-associated remodeling of epithelial identity and tumor-associated extracellular matrix. In contrast, post-treatment samples exhibited increased ER alpha expression, indicating relative enrichment of ER-positive tumor cell populations following neoadjuvant THP (**Fig. 2E**).

### Single-cell profiling reveals ER-associated cancer cell populations linked to treatment resistance

To directly assess intra-tumoral heterogeneity and its association with treatment response, we analyzed 22 baseline breast biopsies using single-cell multiplex CyCIF (**Fig. S1C)** (*14, 18*). Major cell types were annotated, and individual cancer cells were stratified based on HER2, ER, and PR expression, along with Ki-67, PD-L1, and HLA-A (**Fig. S3A**). To ensure accurate quantification of subcellular marker localization, particularly for nuclear-localized ER and PR versus membrane-localized HER2, compartment-specific segmentation masks were applied (**Fig. S3B**).

Single cell analysis revealed stratification of 8 cancer cell populations based on the expression of HER2, ER and PR. ER-positive/HER2-positive tumors displayed a wide range of HER2, ER, and PR expression patterns, while ER-negative/HER2-positive tumors predominantly lacked ER and PR expression, as expected (**Fig. 3A, B**).

**Fig. 3:**
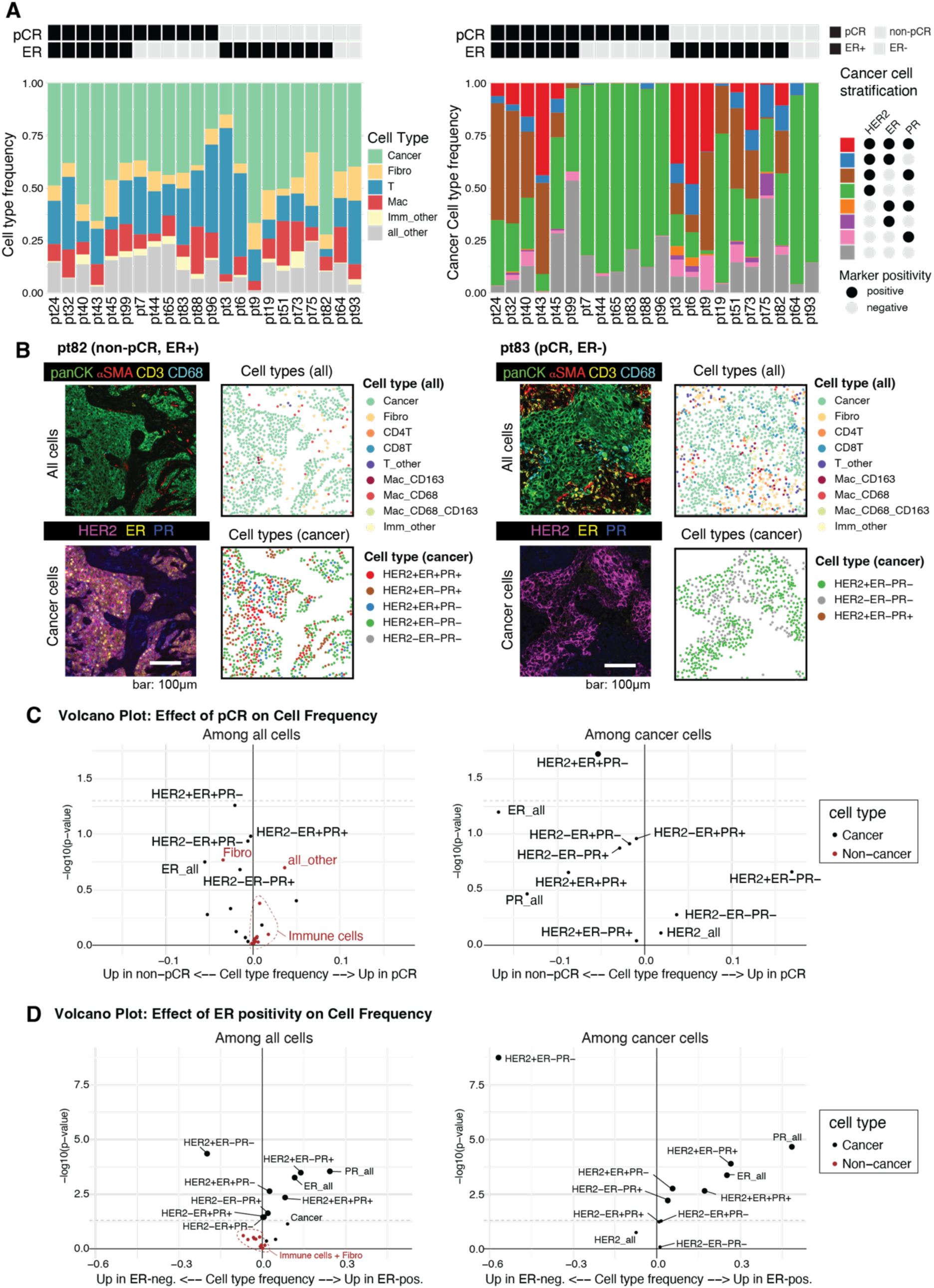
Spatial imaging reveals an association between treatment response and ER-positive cancer cell states. Baseline tumors were assessed using multiplex, single cell CyCIF. **(A)** Frequencies of single cells across patient samples. Left: proportions of major cell types among all cells, shown by pCR and ER status. Right: distribution of cancer cells stratified by HER2, ER, and PR. **(B)** Representative regions of interest (ROIs), showing spatial distribution of cancer and non-cancer cells. Left: CyCIF images; Right: corresponding computationally assigned cell types. **(C)** Volcano plots showing associations between cell frequencies and treatment response. Left panel: frequencies calculated across all cells; Right panel: frequencies calculated among cancer cells. **(D)** Volcano plot showing associations between cell frequencies and clinical ER status.

We assessed whether any of the eight cancer cell populations or immune or stromal cells were associated with neoadjuvant treatment response (pCR vs non-pCR). Of the cancer cell populations, HER2-positive/ER-positive/PR-negative was significantly enriched in non-pCR tumors (**Fig. 3C**). More broadly, the expression of ER on individual cancer cells, regardless of HER2 or PR status, was consistently associated with non-pCR response. To account for the interrelated nature of different cancer cell populations, we applied a LASSO regression model, which confirmed that ER expression (ER-positive/PR-negative/HER2-positive and ER-positive/PR-negative/HER2-negative) remained a strong predictor of non-pCR (**Fig. S4A**). When tumors were stratified by clinical ER status, the expression of ER and/or PR on individual cancer cells was more abundant in ER-positive tumors, while single cells stratified as ER-negative/PR-negative/HER2-positive were enriched in ER-negative tumors, as expected (**Fig. 3D**). In contrast, frequencies of immune cell populations and fibroblasts did not differ significantly by response or ER status, suggesting that bulk immune composition alone does not capture the spatial microenvironmental differences explored in the following section.

### Spatial neighborhood analysis identifies immune exclusion patterns driven by HER2 and ER expression

To test how cancer cell expression of HER2, ER, and PR may influence their spatial context, we conducted a cancer cell–anchored cellular neighborhood (CN) analysis. For each tumor, we defined fixed-radius (20 µm) neighborhoods around cancer cells and randomly sampled 2,000 neighborhoods per specimen for pooled analysis (**Fig. 4A**). Given the marked differences in single cell expression of HER2, ER and PR between the ER-positive and ER-negative tumors (**Fig. 3C**), we stratified the analysis by clinical ER status. All eight cancer cell populations were represented in ER-positive tumors, whereas only two of the populations were present in ER-negative tumors. Accordingly, we focused initial analyses on the 14 ER-positive cases to capture greater phenotypic diversity of the cancer cells. At a high level, we observed substantial diversity in the frequency of major neighboring cell types across different anchor cancer cell populations (**Fig. 4B left, ER-positive**). Among cancer cells, each cancer cell population was most frequently surrounded by neighboring cells of the same type, indicating strong spatial self-association (**Fig. 4B right, ER-positive**).

**Fig. 4:**
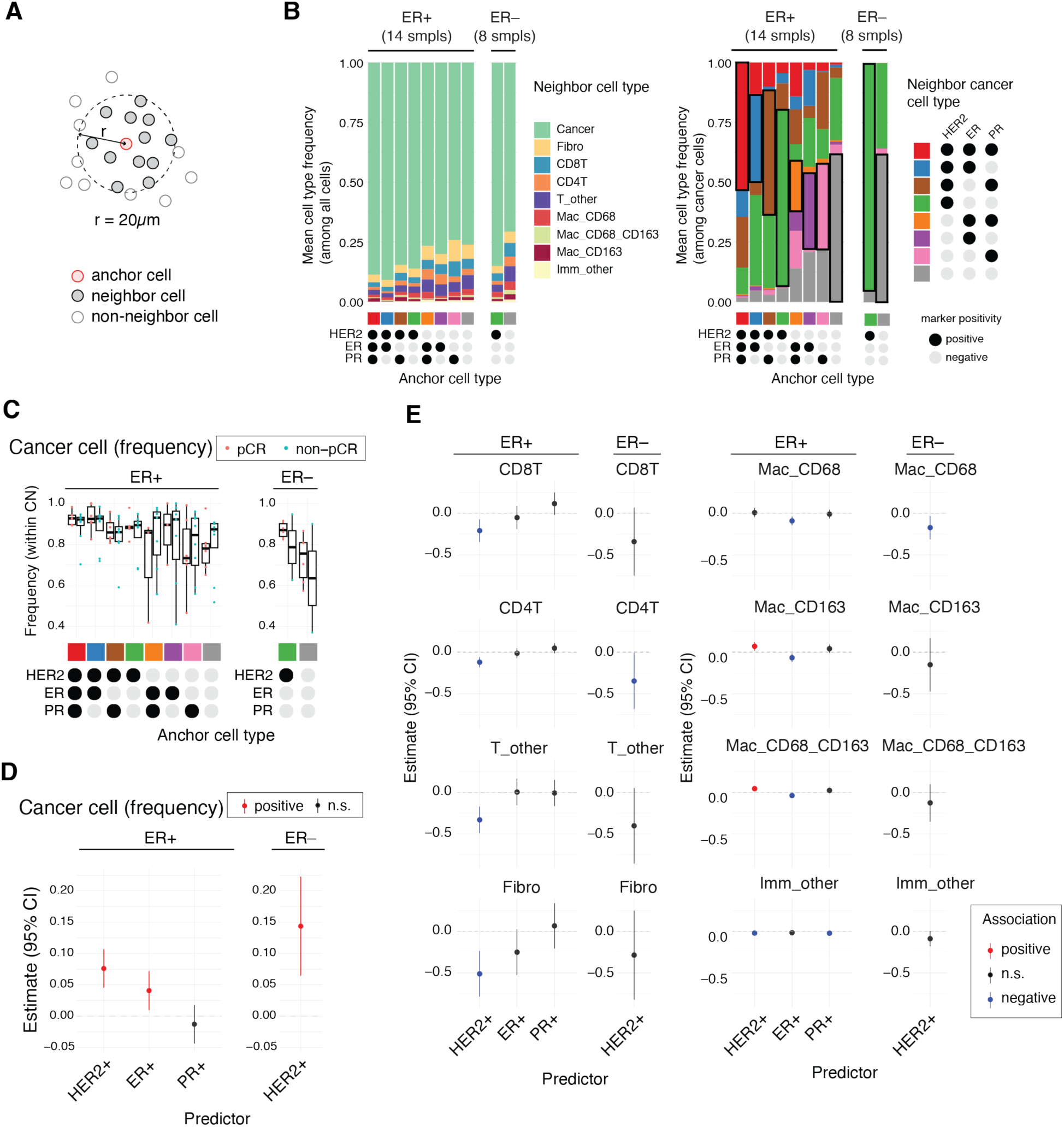
HER2, ER, and PR expression shape the local immune microenvironment of cancer cell neighborhoods. Cellular neighborhood (CN) analysis was performed to investigate how the molecular state of individual cancer cells influences the composition of their surrounding microenvironment. **(A)** Schematic of CN analysis. Each CN was defined as a 20 μm radius surrounding an individual cancer cell (anchor cell). Cells within this radius were designated as neighbor cells; those outside were considered non-neighbors. **(B)** Mean frequencies of neighbor cell types within cancer-anchored CNs, stratified by cancer cell expression of HER2, ER, and PR of the anchor cells. Left: frequencies of cells out of all cell types. Right: frequencies of cancer cells. Black outlines in the right highlight neighboring cancer subtypes that match the anchored cells. **(C)** Mean cancer cell frequency in CNs around 8 cancer cell populations. **(D)** Forest plot showing the association of HER2/ER/PR expression in anchor cells and the frequency of neighboring cancer cells. **(E)** Forest plots showing the association between cancer cell marker expression and the frequency of non-cancer cell types within their neighborhoods. Generalized linear mixed-effects models were fitted separately for ER-positive and ER-negative tumors. Fibro indicates fibroblast cells, Mac indicates macrophages with surface markers as annotated, Imm_other indicates all other immune cells.

Mean cell type frequency across cancer cell populations revealed that both HER2 and ER positivity were significantly associated with increased cancer cell frequency within the neighborhoods, whereas PR positivity did not reveal an association (**Fig. 4C, D**). Because frequency is a relative measure, we also calculated mean cell count per CN, which confirmed the same trend and was adopted as the primary metric for subsequent analyses (**Fig. S4B, C**).

While immune and stromal compositions were broadly similar between pCR and non-pCR tumors, neighborhood composition varied across the eight cancer cell populations (**Fig. S4D**). To investigate whether marker-defined phenotypes influence the local immune environment, we applied linear mixed-effects models. In ER-positive tumors, HER2 expression on individual cancer cells was associated with reduced frequencies of CD8+, CD4+, and other (i.e., CD4- /CD8-) T cell subtypes in their vicinity, suggesting an association with immune exclusion (**Fig. 4E; Fig. S4D, ER-positive**). A similar trend was observed in ER-negative tumors, although statistical power was limited due to smaller sample size (**Fig. 4E; Fig. S4D, ER-negative**).

We further examined the spatial co-occurrence of individual cancer cells. Sample-level analyses confirmed pooled results: anchor cells were typically surrounded by cells with similar expressions of HER2, ER and/or PR (**Fig. S4E**). This pattern was not limited to cells with identical expression of all three markers; cells that matched the anchor cell on two markers were more frequently observed nearby than those matching on only one or none (**Fig. S4F**). Each marker (HER2, ER, PR) contributed additively to the likelihood of neighboring cell abundance (**Fig. S4F**). These findings may reflect gradual phenotypic transitions and local expansion of evolving cancer cell states within each tumor.

We next investigated the expression of HLA-A, PD-L1, and Ki-67 on individual cancer cells. Using generalized linear mixed-effects models, we found that expression of HLA-A and/or PD-L1 on individual cancer cells was positively associated with HER2 and PR expression, but negatively associated with ER expression (**Fig. 5A, B, left and middle panels**). Ki-67 expression was also positively associated with HER2 and negatively associated with ER expression, particularly in ER-negative tumors, consistent with greater proliferative activity in HER2-driven cancer cells. Taken together, these findings suggest that ER expression shapes both the phenotypic and immune architecture of the tumor microenvironment, identifying ER-driven immune exclusion as a candidate mechanism of resistance in HR-positive/HER2-positive breast cancer (**Fig. 5A, B, right panels**).

**Fig. 5:**
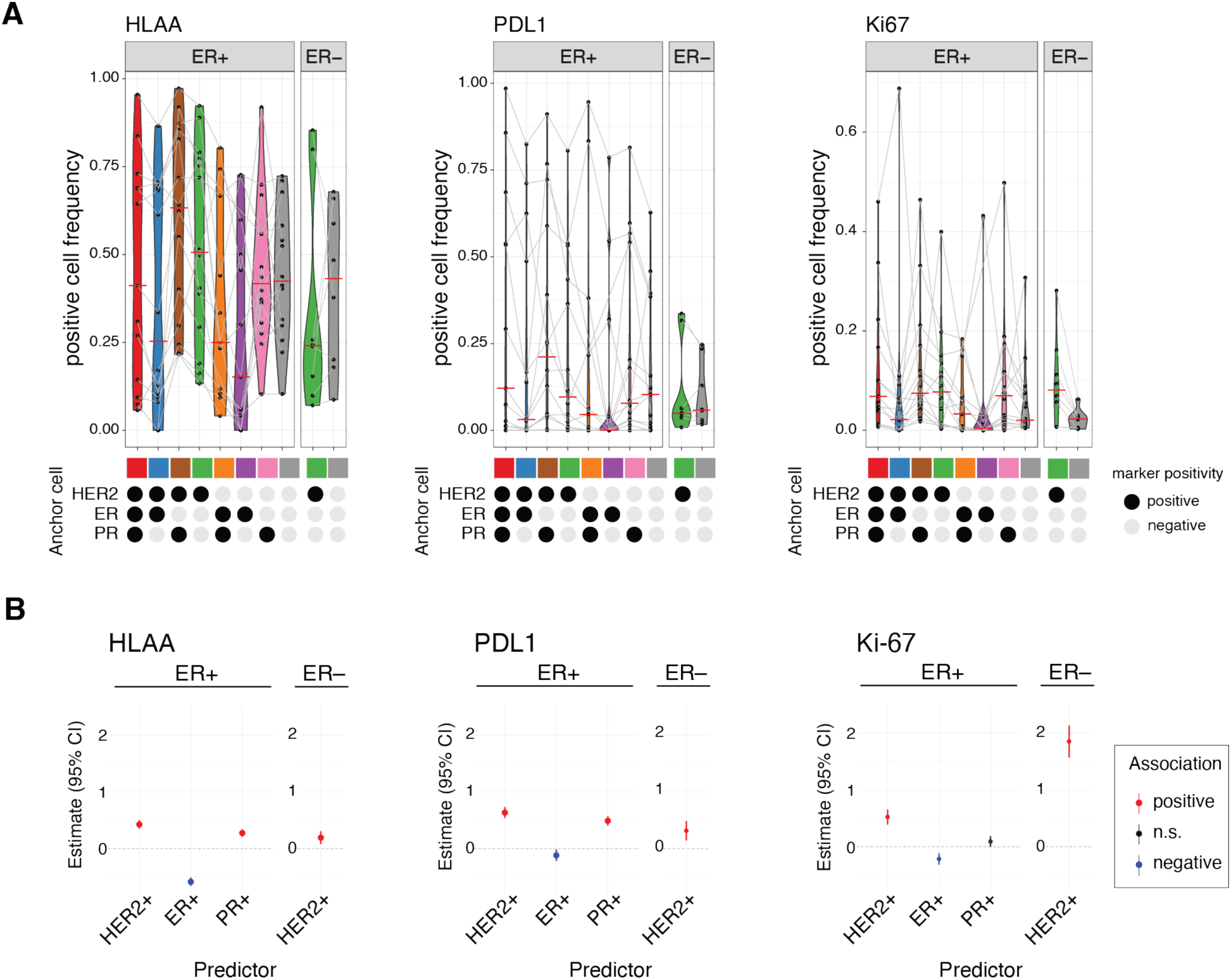
HER2, ER, and PR expression define distinct cancer cell states associated with antigen presentation and proliferation. Baseline tumors were assessed using multiplexed single-cell CyCIF. **(A)** Frequencies of HLA-A, PD-L1, and Ki67 positive cells, stratified by HER2, ER, and PR expression in cancer cells. Each dot represents an individual patient sample; lines connect samples from the same patient. **(B)** Fixed-effect estimates (log-odds ratios) from generalized linear mixed-effects models assessing expression the likelihood of individual cancer cells expressing HER2, ER or PR to express Ki67, HLA-A, or PD-L1.

## DISCUSSION

In this study, we have profiled pre- and post-treatment tumor specimens from patients treated with neoadjuvant THP on the DAPHNe trial, using gene expression, multiplex protein expression, and single-cell spatial analysis. Our work produces a comprehensive overview of the heterogeneity of early-stage HER2-positive breast cancer, and how various features predict treatment response at surgery.

Our findings demonstrate that the strength of HER2 expression is the strongest predictor of favorable response to neoadjuvant THP by both gene (*ERBB2*) and protein (HER2) expression. Numerous studies have demonstrated a similar relationship between HER2 expression and a higher likelihood of pCR, as seen in trials such as NeoALTTO, GeparQuattro, and TRYPHAENA, among others (*19-22*). Previous literature also identifies other surrogates of HER2 gene and protein expression that predict higher likelihood of pCR. For example, among the PAM50 subtypes, the HER2-enriched subtype is most associated with increased pCR likelihood (*23, 24*), whereas higher degrees of intratumor HER2 heterogeneity correlate with lower pCR likelihood (*25*). We further identified ER positivity and its surrogates as the most consistent correlate of lower pCR likelihood by both gene (*ESR1*, *PGR*, ER signaling gene signature) and protein expression, including single-cell spatial analysis. This finding is consistent with prior trials showing ER/PR positivity as a negative predictor of pCR, and is further reinforced here at single-cell resolution (*19, 21, 23, 26*).

Immune-related gene expression signatures, including B cell and T cell signatures, have been established as strong predictive biomarkers of favorable response following neoadjuvant HER2-directed therapy (*22, 27, 28*). The histologic presence of tumor-infiltrating lymphocytes (TILs) has also been shown to be a predictive marker for favorable treatment response in HER2-positive breast cancer (*29, 30*). Interestingly, we did not find any association between treatment response and immune-related gene expression or protein expression (using both GeoMx and CyCIF technologies). This may reflect limited statistical power given the sample size, or insufficient variability in the immune panels used to discriminate response in this cohort.

It is becoming increasingly apparent that HER2-positive/ER-positive and HER2-positive/ER-negative tumors are clinically distinct. In HER2-positive/ER-positive disease, there is complex cross-talk between the two pathways that can affect how the tumor responds to anti-HER2 therapy (*31*). The distinction between ER-positive and ER-negative tumors was apparent in all of our analyses. At the gene expression level, ER-negative tumors had significantly higher expression of the BC p53 and HRD signatures. At the protein expression level, ER-positive tumors exhibited higher expression of the anti-apoptotic protein Bcl-2. In contrast, ER-negative tumors showed increased expression of several immunomodulatory proteins, including PD-L1, CD27, 4-1BB, CD4, and CD20, however, most of these associations were significant only in unadjusted analyses. Notably, despite these protein-level differences, CyCIF analysis did not identify significant differences in immune cell population abundance according to ER status.

Consistent with the higher immune activity characteristic of ER-negative breast cancer (*27, 32*), protein expression analyses by CyCIF and GeoMx revealed substantially more differences between ER-positive and ER-negative tumors than the BC360 gene expression assay. This observation suggests that ER-associated immune variation is driven largely by differences in cellular composition and spatial organization within the tumor microenvironment, which are better captured by spatial proteomic approaches than by bulk gene expression profiling.

Single-cell analyses reveal pronounced intra-tumoral heterogeneity, with coexisting cancer cell subpopulations defined by distinct HER2, ER and PR expression states within individual tumors. We found that cancer cells cluster with other cancer cells with similar expression of HER2, ER, and PR, illustrating local proliferation of tumor cell subpopulations. HER2 expression of individual cancer cells was associated with increased Ki67 expression, particularly in ER-negative tumors, and these HER2-expressing cells were spatially isolated from T cells, suggesting increased proliferation and immune exclusion. Ki67 expression was also negatively associated with ER expression, consistent with the known higher proliferative activity of HER2-positive/ER-negative tumors (*31, 33*). ER expression on individual cancer cells also correlated with reduced HLA-A expression, consistent with prior observations linking ER signaling to diminished antigen presentation in ER-positive breast cancer. These results are consistent with prior work from our group and others demonstrating that ER signaling is inversely correlated with interferon-mediated antigen presentation in ER-positive breast cancer (*34-36*). Higher individual cancer cell expression of ER was also associated with lower expression of PD-L1. Taken together, these findings suggest that HER2 and ER expression not only define cancer cell-intrinsic phenotypes, but may also shape the spatial architecture and immune landscape of the tumor microenvironment, revealing coordinated mechanisms of immune evasion and therapeutic resistance.

Paired comparison of protein expression between baseline and post-treatment samples in patients with unfavorable response (RCB 2/3) revealed remodeling of both tumor cell and stromal immune compartments during neoadjuvant THP. Post-treatment tumors were less proliferative (lower Ki-67 expression), less HER2-driven, and more ER-driven. Similar changes from pre- to post-treatment have been observed in analyses of paired tumor samples from the CALGB 40601, PAMELA, NeoALTTO, and NSABP B-41 trials (*37*). Data regarding immune activation were more mixed, with several immunomodulatory proteins, including PD-L1, CTLA4, FOXP3, CD127, and GZMB decreasing during treatment, and expression of S100B, CD27, 4-1BB, and CD66b increasing during treatment. Current literature is also divided, with some studies finding increased immune activation in residual disease and others finding depletion of immune-related pathways (*37, 38*). The DAPHNe data suggest a nuanced picture of concurrent immune depletion and co-stimulatory enrichment, which warrants further investigation.

This work has several limitations. Each of the biomarker cohorts is slightly different due to distinct tissue requirements and cost limitations, which introduces the potential for heterogeneity in the results. However, characteristics of each cohort are described, and broadly recapitulate the parent trial population. The assessment of tumor changes from pre- to post-treatment are limited by the fact that these were only performed in patients with significant residual disease, due to the difficulty of identifying salient features without tumor cells present.

In conclusion, using complementary gene expression profiling, spatial protein profiling, and multiplex single-cell imaging of prospectively collected specimens from the DAPHNe trial, we demonstrate that HER2 and ER expression are the dominant, reproducible determinants of response to neoadjuvant THP. Importantly, these approaches distinguish heterogeneity across tumors (inter-tumoral) from heterogeneity within tumors (intra-tumoral), revealing how population-level biomarkers and single-cell spatial organization jointly shape therapeutic response. We identify key biologic differences between HR-positive and HR-negative tumors, as well as in baseline vs post-neoadjuvant treatment tumors among patients with significant residual disease at the time of surgery. Single-cell spatial profiling further reveals ER expression on individual cancer cells is associated with reduced local T cell infiltration and HLA-A expression, highlighting the potential contribution of immune exclusion to resistance in HER2-positive/ER-positive tumors. Together, these findings may guide development of more individualized and biomarker-driven treatment strategies.

## MATERIALS AND METHODS

### Study design

DAPHNe (De-escalation to Adjuvant antibodies Post-pCR to Neoadjuvant THP, NCT03716180) was a single-arm phase 2 trial in which 98 patients with stage II-III HER2-positive breast cancer received neoadjuvant paclitaxel (80 mg/m^2^ weekly for 12 weeks), trastuzumab (H; 8 mg/kg initial dose followed by 6 mg/kg for subsequent doses, every 3 weeks for 4 cycles) and pertuzumab (P; 840 mg initial dose followed by 420 mg for subsequent doses, every 3 weeks for 4 cycles). A baseline research biopsy was required for all patients prior to initiation of neoadjuvant therapy. After 12 weeks of treatment, patients underwent surgery. Those who exhibited a pathologic complete response (residual cancer burden [RCB] 0) received adjuvant HP, and those with residual disease (RD) received trastuzumab emtansine (T-DM1). The primary endpoint of the study was adherence to the protocol-specified antibody doublet (without further cytotoxic chemotherapy) in the adjuvant setting for those who exhibited a pCR to determine the feasibility of chemotherapy de-escalation. The primary trial results were previously published (*16*). The Dana-Farber/Harvard Cancer Center Institutional Review Board approved the study and all patients provided written informed consent.

### Gene expression profiling

The nCounter® Breast Cancer 360™ (BC360; Bruker Spatial Biology) gene expression assay was performed on extracted RNA samples using established protocols, as previously described (*39*). This assay measures the expression of 758 genes and 48 curated biological signatures. Briefly, following RNA extraction from formalin-fixed paraffin-embedded (FFPE) tissue specimens, the BC360 assay was used to quantify gene expression levels through direct digital counting of barcoded probes without the need for amplification. Among the evaluated signatures, intrinsic molecular subtypes of breast cancer were determined using the PAM50 gene expression subtypes.

### Proteomic digital spatial profiling

The Bruker Spatial Biology GeoMx® Digital Spatial Profiler (DSP) platform was used to characterize protein expression in both baseline core biopsy specimens and post-treatment surgical samples; the latter were limited to patients with significant residual disease (RCB 2/3). This analysis employed a 58-protein immune cell profiling panel designed to assess the tumor microenvironment, as described in prior publications (*40*). Briefly, tissue sections were first evaluated using hematoxylin and eosin (H&E) staining, which was used to guide the selection of up to 12 regions of interest (ROIs) per slide. Each ROI was further segmented into discrete areas of illumination (AOIs), corresponding to spatially distinct tumor-rich (pan-cytokeratin [pan-CK] positive) regions and stromal (pan-CK negative) regions, as previously described (*41*). Protein targets within each AOI were quantitatively assessed using oligonucleotide-tagged antibodies and UV-photocleavable indexing oligos.

### Multiplexed cyclic immunofluorescence (CyCIF) imaging and spatial analysis

CyCIF was performed as previously described (*14*) to characterize tumor, immune, and stromal cell populations and their spatial organization in pre-treatment breast cancer specimens. Tumor cells were identified using epithelial marker (pan-CK) and subclassified based on estrogen receptor (ER), progesterone receptor (PR), and HER2 expression, while immune and stromal populations were defined using established lineage markers (**Fig. S3A**). Expression of clinically relevant cell-state markers, including antigen presentation and immune checkpoint proteins, was assessed at the single-cell level. CyCIF images were preprocessed and quantified using MCMICRO and OMERO (*42, 43*). Spatial analyses were conducted to assess the cellular composition of local tumor microenvironments by quantifying immune and stromal cells in defined neighborhoods surrounding cancer cells. Generalized linear mixed-effects models accounting for multiple observations per patient were used to evaluate associations between cancer cell marker expression and local microenvironmental composition, with analyses stratified by ER status.

### Cyclic immunofluorescence (CyCIF) imaging

CyCIF was performed as previously described (*14*). Imaging was conducted on 27 pre-treatment breast tumor samples across at least six staining cycles and the images were acquired using CyteFinder II Imaging Instrument (RareCyte). Of 27 samples, five samples were excluded due to severe tissue loss, leaving 22 samples for analysis. The following antibodies were used for the study: PD-L1 (Cell Signaling Technology, #13684), CD3 (Abcam, #ab11089), ER-alpha (Cell Signaling Technology, #74244), HER2 (Abcam, #ab225510), progesterone receptor (Abcam, #ab267523), cytokeratin (pan) (Thermo Fisher Scientific [eBioscience), #41-9003-80/82), androgen receptor (Abcam, #ab194195), alpha-actin-2 (Thermo Fisher Scientific, #41-9760-82), CD45 (BioLegend, #304020, 304056), CD8a (Thermo Fisher Scientific, #53-0008-80), CD4 (R&D Systems, #FAB8165G), CD68 (Cell Signaling Technology, #79594, 79594S), CD57 (BD Biosciences, #561906).

Multiplexed CyCIF images were preprocessed using the MCMICRO pipeline (https://mcmicro.org/) to perform illumination correction, stitching, registration, segmentation, and quantification (*42*). Custom configuration files were deployed to the Harvard Medical School O2 high-performance cluster for batch processing. Quantification was performed across three compartments: cellRing (whole-cell), cytoRing (cytoplasmic/plasma membrane), and nucleiRing (nuclear) (*43*). These data were merged in R based on known marker localization: nuclear (ER, PR, Ki-67), membrane/cytoplasmic (HER2), and whole-cell (CD3, CD4, CD8, CD68, PD-L1, HLA-A, panCK, α-SMA). Final merged tables were exported as .csv files for gating and analysis.

Regions of interest (ROIs) that passed quality control were manually annotated using the OMERO PathViewer and exported via a custom JavaScript utility that queried the OMERO API. Exported ROI coordinates and metadata were used to restrict analyses to high-quality tissue regions. Marker-specific gating thresholds were determined using Gater, a graphical gating tool (https://github.com/labsyspharm/minerva_analysis/tree/gating). Gating was performed manually per sample to define positivity thresholds for each marker. These thresholds were applied to assign binary positive/negative status for downstream cell-type annotation and spatial analysis.

Single-cell quantification data were processed using the CycifAnalyzeR package (https://github.com/kenichi-shimada/CycifAnalyzeR). Cell types were assigned according to marker expression and sample-specific gating thresholds. Clinical metadata, including pCR status and HER2 and ER status determined by CLIA-certified immunohistochemistry, were matched to each sample and used to explore associations with cell-type frequencies, counts, and cell state marker expression. Cell-type frequencies were computed at the sample level and visualized using bar plots stratified by immune, stromal, and tumor compartments. Tumor cells were further subclassified into eight groups based on ER, PR, and HER2 expression. Binary positivity for cell state markers such as HLA-A, PD-L1, and Ki-67 was determined using gated thresholds. The proportion of marker-positive tumor cells was calculated across subtypes and visualized using violin plots to highlight intra-tumor heterogeneity and subgroup differences. Spatial relationships between cells within each sample were analyzed by defining cellular neighborhoods (CNs) based on a 20µm radius around each anchor cell. Neighboring cells within this radius were identified using fixed-radius nearest-neighbor (frNN) algorithms implemented in the computeCN() function from the CycifAnalyzeR package. Up to 2,000 cancer cells per sample were randomly selected as anchors and pooled across samples for downstream analyses.

To assess whether local immune and stromal composition differed by response to therapy, we analyzed frequencies of individual cell types in cancer-anchored CNs stratified by marker-defined cancer cell population (e.g., HER2+ER−PR+). Frequencies were aggregated per sample and compared between pCR and non-pCR groups using unpaired two-sided *t*-tests. Significance levels (*p* < 0.1 and *p* < 0.05) were annotated in boxplots. This analysis revealed minimal differences between pCR groups, prompting us to exclude pCR as a covariate in subsequent modeling. In contrast, ER+ and ER− tumors showed markedly distinct distributions of cancer cell populations. Namely, ER− tumors contained very few ER+ or PR+ cells—justifying separate analyses in ER+ and ER− cohorts to avoid imbalanced comparisons.

### Statistical Methods

#### Modeling associations between cancer cell marker status and neighboring cell type frequencies

To evaluate how tumor-intrinsic biomarker expression relates to microenvironmental composition, we applied linear mixed-effects models using the lme4 package. For each neighboring cell type, we modeled frequency or count as a function of ER, PR, and HER2 positivity in the anchor cancer cell, with sample identity included as a random effect: Frequency or Count ∼ ER+ + PR+ + HER2+ + (1|sample). Models were stratified by ER+ and ER− cohorts. Fixed-effect estimates and 95% confidence intervals were extracted and visualized as dot-and-whisker (forest) plots. Predictors were color-coded by direction and significance: red (positive association, *p* < 0.05), blue (negative), and grey (non-significant). These analyses identified how cancer cell-intrinsic features were associated with enrichment or exclusion of specific immune or stromal cell types in the tumor microenvironment.

#### BC360 and GeoMx DSP Analyses

For all BC360 and GeoMx DSP analyses, in order to control for multiple comparisons, p-values were adjusted using the Benjamini-Hochberg false discovery rate (FDR) approach, with statistical significance defined as an FDR-adjusted p-value <0.05.

#### CyCIF Analyses

Cell-type frequency within cancer-cell-anchored CNs was averaged across all qualifying CNs (per sample, per anchor-cell subtype) to give one value per sample. Associations between this sample-level frequency and pCR, and separately with ER status, were each assessed with an unpaired two-sided Welch’s t-test. Associations with tumor-intrinsic biomarker status were modeled with linear mixed-effects models (lme4/lmerTest; Gaussian, cell-type frequency as the outcome) with a random intercept for sample to account for multiple CN-anchor subtypes per patient: for ER+ tumors, frequency ∼ ER + PR + HER2 status (of the anchor-cell cancer subtype) + (1 | sample); for ER− tumors, restricted to the two HER2-defined subtypes (i.e., HER2+/ER– /PR– or HER2–/ER–/PR–) present in that stratum, frequency ∼ HER2 status + (1 | sample). All tumor-intrinsic-biomarker analyses were stratified by ER status because cancer-cell subtype composition differed substantially between ER+ and ER− tumors.

Separately, whether a neighboring cancer cell expressed a given marker (HER2, ER, or PR; binary positive/negative call) was modeled as a function of the CN anchor cell’s tumor-intrinsic biomarker status using a generalized linear mixed-effects logistic model (lme4, binomial family), with a random intercept for sample. Cells were downsampled to at most 2,000 cancer cells per sample (fixed seed) before fitting. For ER+ tumors: expression ∼ HER2 + ER + PR + (1 | sample); for ER− tumors: expression ∼ HER2 + (1 | sample).

p-values were not adjusted for multiple comparisons; significance was assessed at nominal p < 0.05 throughout, including the volcano plots (Fig. 1E,F) and the mixed-model and dot-whisker/forest-plot summaries across cell types.

## List of Supplementary Materials

Fig. S1 to S4

Tables S1 and S2

## Acknowledgments

The authors wish to acknowledge Kaitlyn Bifolck, BA, for manuscript editing and submission assistance. She is a full-time employee of Dana-Farber Cancer Institute.

## Author contributions

Conceptualization: EC, KS, EPW, JG, EAM, AGW

Data curation: EC, IS, KS, TL, ERO, FNN, ERM, NTayob, JG, AGW

Formal analysis: EC, IS, KS, TL, JD, J-RL, AS, MGT, KZ, CWW, RP, FNN, ERM, NTayob, JG, AGW

Funding acquisition: AGW, EPW, EAM

Investigation: EC, IS, KS, TL, JD, J-RL, AS, MGT, KZ, CWW, RP, FNN, ERM, NS, LMS, MF, MC, ND, NTung, NTayob, JG, AGW

Methodology: EC, IS, KS, TL, JG, EAM, AGW

Project Administration: ERO, AP Resources: N/A

Software: KS, FNN, ERM, JG

Supervision: EPW, JG, EAM, AGW

Validation: KS, TL, MGT, FNN, ERM, NTayob, JG

Visualization: EC, IS, KS, JG, AGW

Writing – original draft: EC, IS, KS, AGW

Writing – review & editing: All authors

## Data and materials availability

The compiled CyCIF data, including cell-level expression tables, ROI metadata, gating thresholds, and cell-type annotations, will be made publicly available at the time of peer-reviewed publication, as a deliberate embargo pending publication of the primary manuscript. Code used for image preprocessing, quantification, gating, and spatial analysis is available at https://github.com/GuerrieroLab/DAPHNe.

