## Supplementary material for "Spatial/molecular heterogeneity and treatment response in HER2+ early breast cancer: Translational analyses from the DAPHNe trial"

**Tables S1 and S2**

**Figs. S1 to S4**

**Table S1:** Protein expression differences between hormone receptor positive (HR+) and hormone receptor negative (HR-) tumors

| | Stroma compartment | Tumor compartment |
|---|---|---|
| Higher in HR+ | ER alpha* (FC 2.25) | ER alpha* (FC 8.94) |
| | Fibronectin (1.68) | Bcl-2* (1.98) |
| | B7-H3 (1.44) | Fibronectin* (1.90) |
| | | PR (1.69) |
| | | TIM-3 (1.37) |
| Higher in HR- | CD27* (FC 1.36) | IDO1 (FC 1.50) |
| | IDO1 (1.77) | S100B (1.44) |
| | CD45 (1.45) | MART1 (1.33) |
| | CD3 (1.35) | |
| | CD20 (1.34) | |
| | STING (1.33) | |
| | CD4 (1.30) | |
| | ICOS (1.30) | |
| | CD45RO (1.28) | |
| | CD40 (1.28) | |
| | CD127 (1.27) | |
| | VISTA (1.25) | |
| | PD-L1 (1.24) | |
| | 4-1BB (1.21) | |

Parenthetical number after each protein indicates fold-change compared to other group (eg, HR+ compared to HR-, or HR- compared to HR+).
Abbreviations: FC = fold-change, HR+ = hormone receptor positive, HR- = hormone receptor negative
*significant at FDR p-value <0.05 (If no asterisk indicated, the result is significant at p-value <0.05 without FDR correction.)

**Table S2:** Protein expression predictors of residual cancer burden (RCB) response

| Overall population | | |
|---|---|---|
| Higher in RCB 0/1 | HER2* (FC 3.75) | |
| | PD-L1 (1.25) | |
| Higher in RCB 2/3 | ER alpha* (FC 2.44) | |
| | BCL-2 (1.46) | |
| HR+/HER2+ patients | | |
| | Stroma compartment | Tumor compartment |
| Higher in RCB 0/1 | HER2* (FC 4.28) | HER2* (FC 4.21) |
| | | CTLA4 (1.64) |
| Higher in RCB 2/3 | | Bcl-2* (2.18) |

The HR-/HER2+ subgroup was not evaluated given that the number of HR-negative patients with RCB 2/3 response (N = 2) was too small to make valid comparisons.
Parenthetical number after each protein indicates fold change compared to other group (eg, RCB 0/1 compared to RCB 2/3).
Abbreviations: FC = fold change, RCB = residual cancer burden, HR+ = hormone receptor-positive
*significant at FDR p-value <0.05 (If no asterisk indicated, the result is significant at p-value <0.05 without FDR correction.)

**Fig. S1A:** REMARK diagram for Nanostring BC360 gene expression analysis.

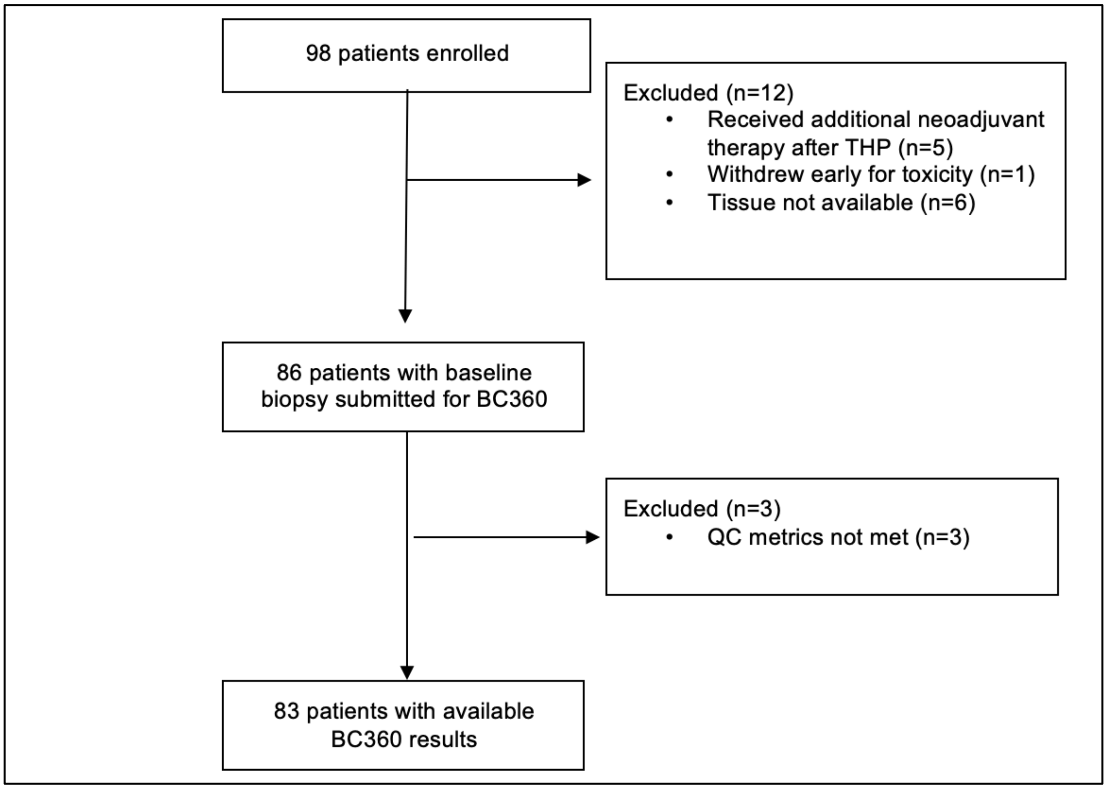

Abbreviations: THP, paclitaxel/trastuzumab/pertuzumab; QC, quality control.

**Fig. S1B:** REMARK diagram for Nanostring GeoMx protein expression analysis.

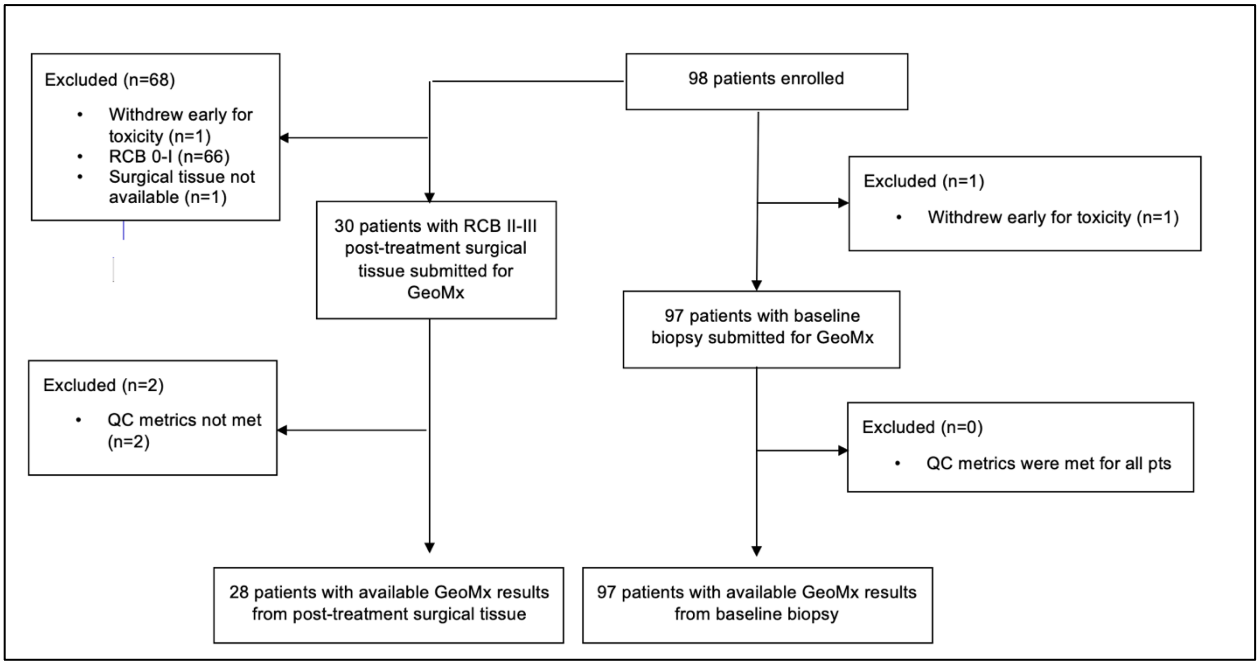

Abbreviations: THP, paclitaxel/trastuzumab/pertuzumab; QC, quality control; RCB, residual càncer burden.

**Fig. S1C:** REMARK diagram and patient characteristics for CyCIF analysis (n=22)

CyCIF patient characteristics:

| | **CyCIF sub-population** | |
|---|---|---|
| **Age, years (median and range)** | 47 (33-76) | |
| **Clinical tumor stage** | | |
| cT1 | 4 | 18.2% |
| cT2-3 | 18 | 81.8% |
| **Clinical nodal stage** | | |
| cN0 | 14 | 63.6% |
| cN1-3 | 8 | 36.4% |
| **Pathological response** | | |
| RCB 0 | 12 | 54.5% |
| RCB I | 1 | 4.5% |
| RCB II | 8 | 36.4% |
| RCB III | 1 | 4.5% |
| **Estrogen receptor status** | | |
| Positive* | 14 | 63.6% |
| High-positive | 14 | 63.6% |
| Low-positive | 0 | 0% |
| Negative | 8 | 36.4% |

*ER-negative is defined as ER 0%. ER high-positive defined as ER or PR $\geq$10%. ER low-positive defined as everything else.

CyCIF REMARK:

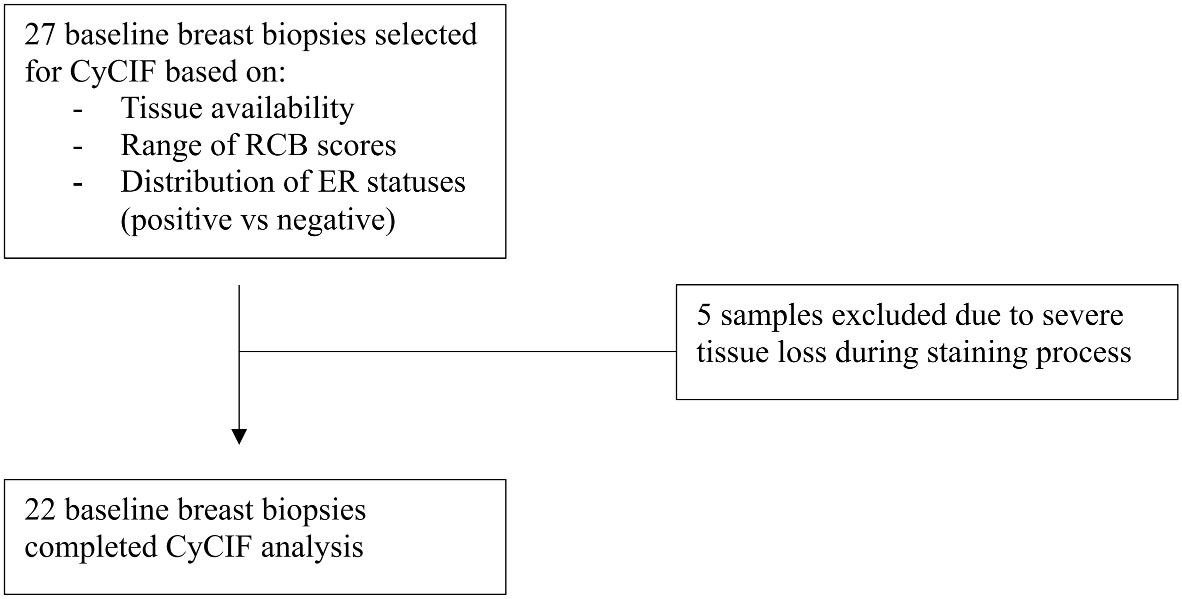

**Fig. S2:** Gene expression differences by nodal status

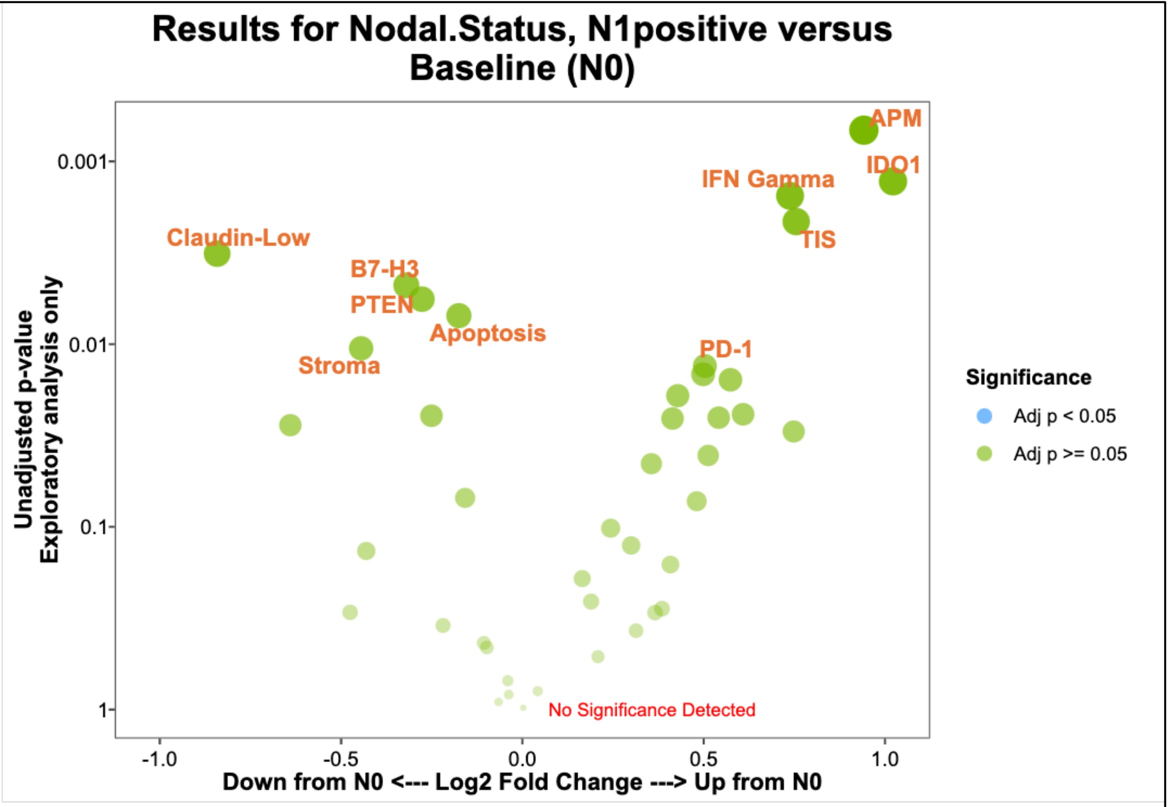

**Fig. S3:** Spatial imaging reveals an association between treatment response and ER+ cancer cells, but not with immune or stromal cells. **(A)** Cell type classification based on marker expression. Cells were assigned to major cell types (cancer, immune, or stromal) based on lineage marker expression. Cancer cells were further stratified according to HER2, ER, and PR status. **(B)** Examples of protein expression using nuclear and non-nuclear masks. Representative examples of HER2 and ER show distinct subcellular localization patterns. Blue outlines denote optimal masks used to measure nuclear and non-nuclear expression in single cancer cells.

### Figure S3

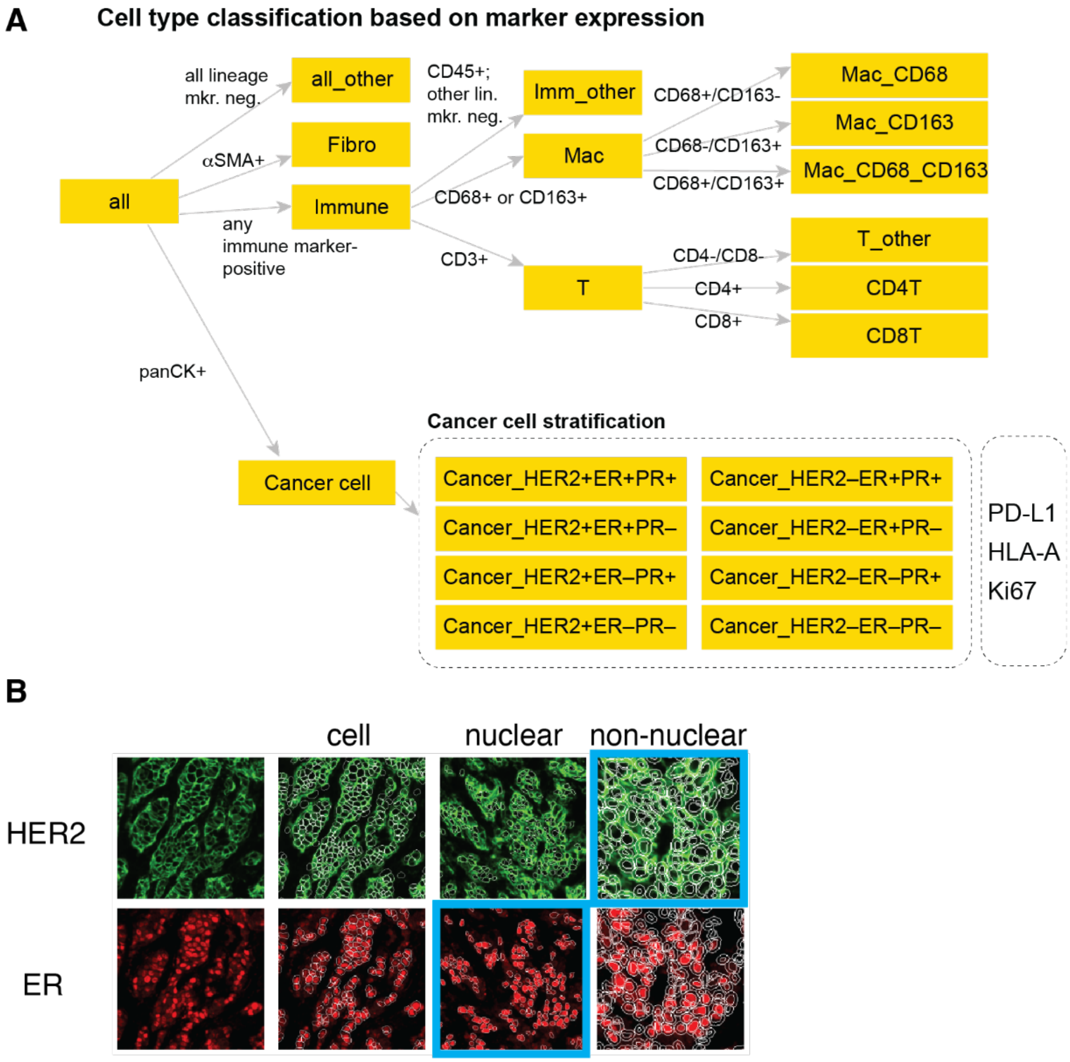

**Fig. S4:** Spatial distribution of immune and cancer cells within cancer cell-centered cellular neighborhoods (CNs). **(A)** LASSO coefficient paths showing the relative association of cell abundance with pCR. **(B)** Mean count of neighboring cancer cells within CNs around cancer-centered CNs, stratified by cancer cell expression of HER2, ER, and PR of the anchor cells. **(C)** Forest plots showing the association between cancer cell marker expression and the count of non-cancer cell types within cancer-centered CNs. **(D)** Box plots showing the frequency of immune and stromal cell types within cancer-centered CNs, grouped by expression of HER2, ER, and PR of individual cancer cells. Each dot represents a patient sample. Frequencies reflect the proportion of neighboring immune/stromal cells surrounding anchor cancer cells. Data are stratified by pCR status and computed separately for ER-positive and ER-negative samples. Asterisks (* and **) indicate differences between pCR and non-pCR groups with $p < 0.1$ and $p < 0.05$, respectively. **(E)** Box plots showing the frequency of neighboring cancer cells within cancer cell-centered CNs, grouped by expression of HER2, ER, and PR of individual cancer cells. As in (A), anchor cells are stratified by HER2, ER, and PR status. Highlighted rectangles indicate subtypes where the marker status of neighboring cancer cells matches that of the anchor cells, reflecting maximal local enrichment. **(F)** Forest plots showing the effect of the expression of HER2, ER, and PR of individual cancer cells on the frequency of neighboring cancer cells. Each estimate reflects the association between marker expression in anchor cells and the abundance of matching cancer cells within their CNs. Analyses are stratified by ER status.

### Figure S4

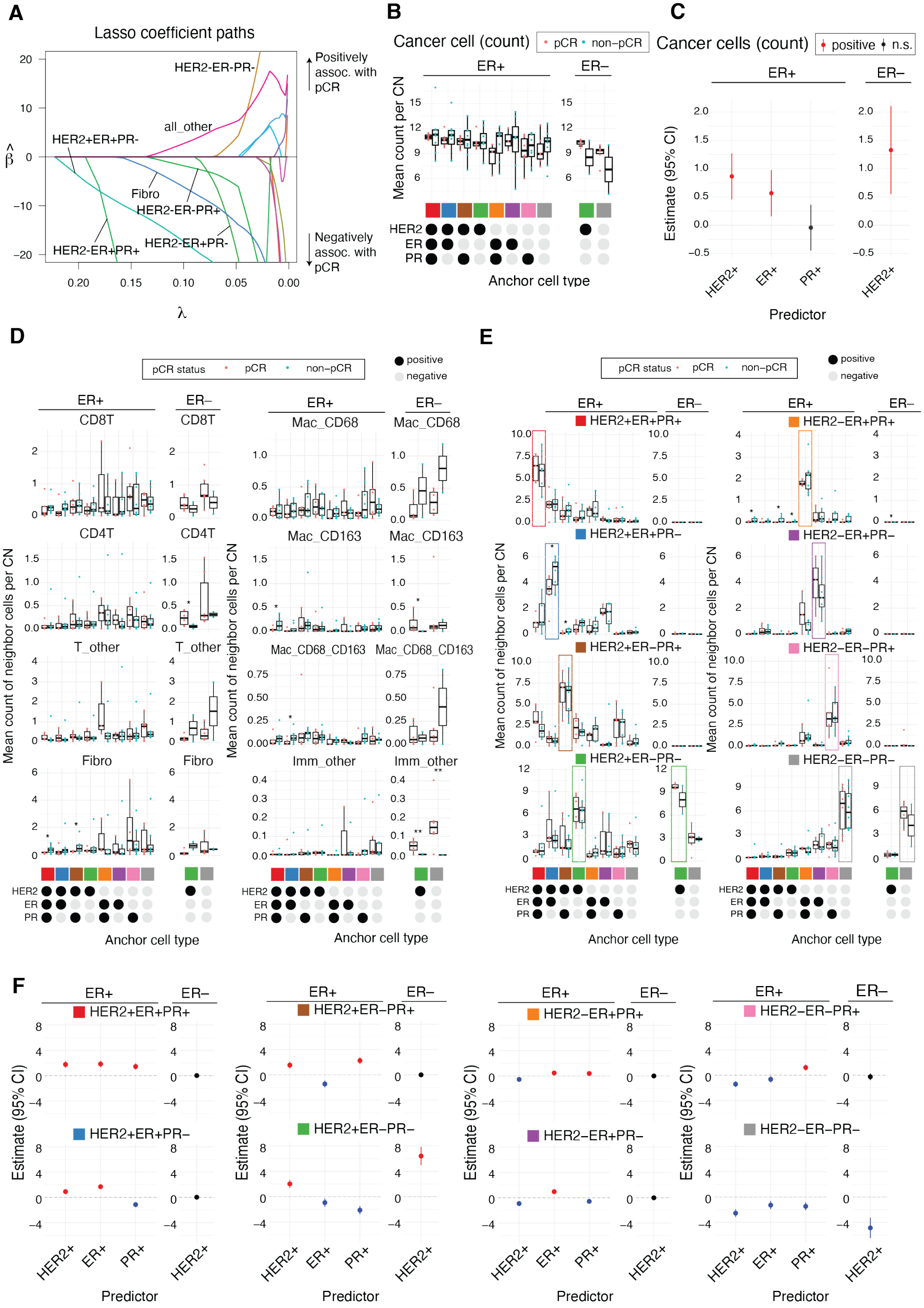

A
Lasso coefficient paths
Positively assoc. with pCR
Negatively assoc. with pCR
HER2-ER-PR-
all_other
HER2+ER+PR-
Fibro
HER2-ER-PR+
HER2-ER+PR+
HER2-ER+PR-
λ
B
Cancer cell (count)
pCR non−pCR
ER+
ER−
Mean count per CN
Anchor cell type
C
Cancer cells (count)
positive n.s.
Estimate (95% CI)
Predictor
D
pCR status pCR non−pCR
positive negative
CD8T
CD4T
T_other
Fibro
Mac_CD68
Mac_CD163
Mac_CD68_CD163
Imm_other
Mean count of neighbor cells per CN
Anchor cell type
E
HER2+ER+PR+
HER2+ER+PR−
HER2+ER−PR+
HER2+ER−PR−
HER2−ER+PR+
HER2−ER+PR−
HER2−ER−PR+
HER2−ER−PR−
F
Estimate (95% CI)
Predictor
